# Rapid fluctuations in histamine associated with intake of nutritive and non-nutritive solutions

**DOI:** 10.1101/2024.11.07.622425

**Authors:** KL Volcko, A Gresch, B Benowitz, H Taghipourbibalan, M Visser, GD Stuber, AG Gordon-Fennell, T Patriarchi, JE McCutcheon

## Abstract

The neurotransmitter histamine is involved in control of food intake, yet its dynamics during individual feeding episodes remain unexplored. Therefore, we used the novel genetically-encoded histamine sensor, HisLightG, combined with fiber photometry to measure histamine release in two hypothalamic regions critical for the food-suppressive effects of histamine, the paraventricular nucleus of the hypothalamus (PVH), and the ventromedial hypothalamus (VMH). Male mice were tested under different conditions to assess whether hunger, time of day, or the caloric content of the solution they were given affected histamine fluctuations. We found that histamine levels changed rapidly in response to eating. These histamine fluctuations were influenced by experimental conditions, with slightly smaller responses when the test solution was sucralose (both regions) or during the light cycle (PVH only). Notable regional differences were identified, such that in the PVH histamine rebounded to baseline levels, whereas in the VMH histamine remained lower than baseline for at least 10 seconds after licking ceased. In a separate cohort of male and female mice, enhancing histamine tone via administration of a histamine precursor (L-histidine) reduced the number of licks across multiple sucrose concentrations. Together, these findings indicate that histaminergic activity is modulated rapidly during ingestive episodes, and that understanding these release patterns will give insight into histamine’s role in appetite suppression.

## 1. Introduction

The mechanisms that govern meal termination and reductions in food intake are complex and intertwined. The neurotransmitter and neuromodulator histamine plays an important role in ingestive behavior and many well-known satiety signals, including leptin (Morimoto et al., 1999; Toftegaard et al., 2003), amylin (Lutz et al., 1996; Mollet et al., 2003), cholecystokinin (Attoub et al., 2001), and glucagon-like peptide-1 (Gotoh et al., 2005), require histamine in order to exert their effects on eating behavior. Histamine may be responding directly to some of these signals – for example, amylin fibers and histaminergic neurons are found in close proximity (D’Este et al., 2001) – but there is likely also indirect input, such as through the melanocortin system (Michael et al., 2020) or noradrenaline (Kurose & Terashima, 1999).

Histamine is involved in many physiological processes besides eating, and can either stimulate or suppress food intake depending on the precise location and receptor subtype activated. The bulk of previous research measuring histamine levels in the brain used microdialysis, a technique which is limited by a relatively large sampling area (e.g., “anterior hypothalamus”) as well as low temporal resolution (e.g. 20-30 min between samples). Fast-scan cyclic voltammetry has also been used to measure histamine with high temporal resolution *in vivo* (Berger et al., 2022; Samaranayake et al., 2015). However, properties of the molecule and its release patterns make these studies difficult to perform and have limited utility for studying spontaneous histamine release during naturally occurring behaviors (i.e., outside of stimulation of histamine containing fibers). Recent advances in genetically-encoded fluorescent neurotransmitter sensors have allowed *in vivo* neurotransmitter release to be combined with fiber photometry yielding measurements with high chemical, temporal and spatial precision (Patriarchi et al., 2019; Simpson et al., 2024). So far, a histamine sensor, GRAB-HA, has been used in two other studies showing fluctuations during sleep-wakefulness (Dong et al., 2023) and after an aversive stimulus (Lin et al., 2023). To our knowledge, however, no one has previously recorded histamine fluctuations using fiber photometry during eating behavior. Moreover, unlike the other studies cited, we use the novel histamine sensor HisLightG (Kagiampaki et al., 2023).

Central histamine-producing neurons are located in the tuberomammillary nucleus in the posterior hypothalamus (Panula et al., 1984; Watanabe et al., 1984), and project widely throughout the brain, including many hypothalamic nuclei (Lin et al., 2023; Panula et al., 1989). Histamine acts on several receptor subtypes, although it is the H1 receptor (H1R) that mediates suppressive effects of histamine on food intake (Fukagawa et al., 1989; Kurose & Terashima, 1999; Lecklin et al., 1998; Masaki et al., 2004; Morimoto et al., 1999; Ookuma et al., 1989; Sakata et al., 1988). H1Rs are expressed in many parts of the brain, with a high density found in both the ventromedial hypothalamus (VHM) and the paraventricular nucleus of the hypothalamus (PVH) (Futagawa et al., 2024). Infusions of H1R antagonists, or drugs to inhibit local histamine production, into specific hypothalamic nuclei have indicated that these two regions – the VMH and the PVH – are the critical sites of action for histamine to attenuate eating (Ookuma et al., 1989, 1993; Sakata et al., 1988, 1990, 1991).

We wanted to examine the patterns of histamine release in the PVH and the VMH during eating bouts in mice, using fiber photometry and the new histamine sensor, HisLightG (Kagiampaki et al., 2023). Histamine levels in the hypothalamus are sensitive to time of day (Doi et al., 1994; Mochizuki et al., 1992; Sakata et al., 1988). We therefore recorded histamine fluctuations during both the dark and light phase. Additionally, histamine levels may or may not be influenced by hunger status (Itoh et al., 1991; Sakata et al., 1994). Consequently, we measured histamine in both *ad libitum* and food restricted conditions. Finally, in a separate experiment to determine how histamine levels shape consumption over a range of rewarding solutions, we manipulated histamine levels by providing the precursor, L-histidine, and measured licking for different concentrations of sucrose.

## 2. Methods

### 2.1 In vitro experiments

#### 2.1.1 Cell culture, confocal imaging and quantification

HEK293T cells (ATCC #CRL-3216) were cultured in DMEM medium (Thermo Fisher) supplemented with 10% FBS (Thermo Fisher) and 100 μg/ml Penicillin-Streptomycin mix (Thermo Fisher) and incubated at 37°C with 5% CO_2_. Cells were transfected at 50-60% confluency in glass-bottomed dishes using PolyFect Transfection Reagent (Qiagen) according to manufacturer instructions, and used for follow-up experiments 24–48 h after transfection. Before imaging, cells were rinsed with 1 mL of Hank’s Balanced Salt Solution (HBSS, Life Technologies) containing CaCl_2_ and MgCl_2_ supplemented with 30 mM HEPES, both from Thermo Fisher. All cells were imaged at room temperature in glass bottom dishes using HBSS with 30 mM HEPES. Imaging was performed using Zen Blue software on an inverted Zeiss LSM 800 confocal microscope equipped with 488 nm laser and a 40X oil-based objective. During timelapse imaging, ligands were applied manually using a micropipette to reach a final concentration of 10 μM. For quantification of the fluorescence response ΔF/F_0_, regions of interest (ROI) enclosing isolated cell membranes were selected manually using the threshold function of Fiji (ImageJ). Sensor response (ΔF/F_0_) was calculated as follows: (F(t)[−[F_0_)/F_0_ with F(t) being the ROI mean gray value at each time point (t), and F_0_ being the mean gray value of the ten timepoints immediately prior to ligand addition.

### 2.2 Photometry Experiments

#### 2.2.1 Animals

Adult male C57BL/6NRj mice (6 – 8 weeks old) were purchased from Janvier (France) and housed in a temperature- and humidity-controlled room, maintained on a 12:12 light:dark cycle (lights-on at 00:00). Two mice shared each cage, separated by a perforated divider that allowed visual, olfactory, and auditory communication but prevented major physical contact. Water and food were available *ad libitum* unless mice were undergoing food restriction. All animal care and experimentation followed the EU directive 2010/63/EU for animal experiments, and was approved by the National Animal Research Authority in Norway.

#### 2.2.2 Surgical procedures

Surgery was performed after a minimum of 5 days of habituation to the animal facility. Mice (n = 23) were anesthetized with isoflurane and secured in a stereotaxic apparatus (Kopf, Tujunga, CA, USA). Each mouse was given subcutaneous injections of buprenorphine (0.1 mg/kg; Temgesic) and meloxicam (5 mg/kg; Metacam) to provide systemic analgesia, as well as bupivacaine (1 mg/kg; Marcain) as a local anesthetic. A small burr hole was made above the target area (see Table 1 for coordinates; all coordinates based on a flat skull) and 500 nl (0.6 x 10^9^ vg) of AAV-9/2-hSyn1-chI-HisLightG-WPRE-bGHp(A) (#10264; ETH Viral Vector Facility) was infused at a rate of 100 nl / minute using a Nanofil syringe with 33 g blunt needle and UMP3 microsyringe pump (WPI; Sarasota, FL, USA). The needle remained in place for 10 min to allow the virus to diffuse. Next, a fiber optic cannula (1.25 mm ceramic ferrule, 0.4 NA, 400 µm core; RWD) was lowered slowly to just above the virus injection. The fiber optic was secured to the skull with dental cement (Superbond C&B). Each mouse received meloxicam 2 days postoperatively.

**Table 1.** Surgical coordinates. Coordinates used to target the PVH and VMH are provided, including the medial-lateral, anterior-posterior, and dorsal-ventral coordinates relative to bregma and based on a flat skull.

| Target | Medial-Lateral | Anterior-Posterior | Dorsal-Ventral (virus) | Dorsal-Ventral (fiber) |
| --- | --- | --- | --- | --- |
| Paraventricular nucleus of the hypothalamus | 0.3 mm lateral to midline | 0.8 mm posterior to bregma | 4.70 mm ventral to skull | 4.60 mm ventral to skull |
| Ventromedial nucleus | 0.3 mm | 1.20 mm | 5.65 mm ventral | 5.55 mm ventral |
| of the hypothalamus | lateral to<br>midline | posterior to<br>bregma | to skull | to skull |

#### 2.2.3 Testing apparatus

Testing took place in operant chambers (24 cm x 20 cm; Med Associates, Fairfax, VT). The fan was on continuously during testing. When mice were tested during their dark phase, the house light was off in the testing chambers, and when the mice were tested during their light phase, the house light was on in the operant chambers. Solutions were delivered through blunted 18 g needles attached to 5 ml syringes via flexible tubing. These syringes were mounted on syringe pumps that dispensed the solution when licks were detected via a contact lickometer. These pumps were controlled by MEDPC-V software (Med Associates).

While the mice freely explored the operant chambers and licked solutions, histamine activity was recorded via fiber photometry. An RZ10x system and Synapse software (Tucker-Davis Technologies, Alachua, FL, USA) controlled two LED light sources, one blue (465 nm) and one violet (405 nm). The blue light was sinusoidally modulated at 330 Hz and the violet at 210 Hz. A fluorescence minicube (Doric Lenses, Quebec, Canada) combined the blue and violet wavelengths. A 400 µm optical patch cord (Doric Lenses) transmitted the light to the mouse, where the cable and ferrule were attached by a black ceramic sleeve (RWD). This same cable was used to collect light emitted from the brains of the mice. This light was detected by a femtowatt photoreceiver (#2151; Newport, Irvine, CA, USA).

#### 2.2.4 Testing procedures

Each mouse underwent a series of testing sessions in different blocks, with each block a different condition. The order of the blocks was the same for all mice. Testing began at least 12 days after surgery and after mice had been habituated to having a cable attached to their fiber optic implant. The four blocks were 1) *ad libitum* light, 2) food restricted dark, 3) *ad libitum* dark, and 4) *ad libitum* sucralose. In the *ad libitum* light block (“Light”), mice with *ad libitum* food and water were tested during the light phase. They were placed in operant chambers for three 30-minute sessions on different days, in which 1 kcal / ml “Ensure” (Nestle Resource Complete neutral flavor) was continuously available through a spout. In the food restricted dark block (“Restricted”), mice were tested in the dark while food restricted. Food restriction was achieved by giving 2.5 – 3 g of chow to each mouse each day; this resulted in approximately 5-10 % weight loss. In this phase they were again given “Ensure” during three 30 min sessions. The *ad libitum* dark block (“Dark”) was identical to the Light block except mice were tested during their dark phase and only tested in 2 sessions. Lastly, in the *ad libitum* sucralose block (“Sucralose”) mice were tested without food restriction in the dark, but given 2 mM sucralose in the single 30 min session.

#### 2.2.5 Histology

Only mice with virus and fibers placed correctly were included in the final analysis. To check placements, brains were collected at the end of the experiment. Briefly, mice were deeply anesthetized with 0.3 ml of ZRF mix (zolazepam, 3.3 mg/ml; tiletamine, 3.3 mg/ml; xylazine, 0.45 mg/ml; fentanyl, 2.6 ug/ml). The mice were then transcardially perfused with heparinized saline and 4% paraformaldehyde. Brains were removed, postfixed overnight in 4% paraformaldehyde, and placed in 30% sucrose with Proclin. A microtome (Leica, Deer Park, IL; USA) was used to cut brains into 40 µm coronal sections. Every third section was directly mounted onto a Superfrost Plus slide and coverslipped. The native fluorescence of the viral injections was used to determine the location and spread of the injection, and the fiber tract used to assess location of the implant.

### 2.3 Histamine manipulation experiments

#### 2.3.1 Animals

Adult male and female C57BL/6J mice (n=4 of each sex) were bred and housed in the animal facility at University of Washington. Mice were kept on reverse 12h light:dark cycle (lights-on at 21:00) and behavioral experiments were conducted within the dark cycle. Mice were single-housed to prevent damage to the head fixation device, provided with water *ad libitum*, and during experiments were food restricted to ∼90% of free-feeding weight by providing a ration of food (2.5-4 g / day) depending on weight change from the previous day. All animal procedures were pre-approved by the Institutional Animal Care and Use Committee (IACUC) at University of Washington (protocol 4450-01).

#### 2.3.2 Surgery

Mice (n=8) were anesthetized with isoflurane (5% for induction, 1.5 – 2% for maintenance), injected with analgesic (carprofen, 10 mg/kg, s.c.), and the scalp was shaved using electric clippers. Mice were mounted in a stereotaxic frame with heat support and the scalp was injected with local anesthetic (lidocaine, 2%, s.c.) and sterilized using ethanol and betadine. An incision was made using a scalpel and the skull was cleared of tissue and scored with the pointed end of the scalpel. The skull was leveled and 2 micro screws were inserted into burr holes drilled in the lateral portion of the occipital bone, with one additional screw placed in the rostral portion of the skull. A head fixation device (a head plate; 0.8 mm stainless steel 304 series, sendcutsend.com) was attached the skull and screws using Super Glue and then dental cement was used to encase a portion of the head bar and the screws. Once fully set, mice were removed from the stereotaxic frame, allowed to recover with heat support, and then returned to their home cage and allowed to recover for at least 1 week before food restriction began.

#### 2.3.3 Behavior in multispout apparatus

Habituation and training took place as described in (Gordon-Fennell et al., 2023). Briefly, mice were first habituated to gentle scruffing, handling by their head bar, and light restraint within a 50 mL conical over 3 days. Next, mice were given free access to sucrose when head-fixed in the multispout apparatus for 10 min on a single day. Mice were then trained on the multispout apparatus with multiple concentrations of sucrose available. The assay consisted of daily sessions with 100 trials with 3 s access to one of five different sucrose concentrations (0% (water), 5%, 10%, 20%, and 30%, 20 trials of each). Solutions were presented pseudorandomly with an inter-trial interval of 12.5 ± 2.5 s. Licks were detected using a capacitive touch sensor (Adafruit MPR121). Once mice had reached stable behavior in which lick rate showed a monotonic relationship with sucrose concentration, the pharmacological manipulation began. In 2 consecutive sessions mice received either L-histidine (500 mg/kg) or vehicle (saline) i.p. 2 h before being tested in the multispout apparatus. Order of administration was counterbalanced across mice and sexes.

### 2.4 Experimental Design and Statistical Analysis

#### 2.4.1 In vitro experiments

*In vitro* experiments used Welch ANOVA with Dunnett’s multiple comparison test. Each neurotransmitter was tested on 30 cells. All experiments were repeated 3 times with similar results. A p-value of < 0.05 was considered significant.

#### 2.4.2 Photometry experiments

For the photometry experiment, all statistical analyses were conducted using JASP (https://jasp-stats.org/). Post hoc tests were Holm corrected for multiple comparisons. Data from mice in the PVH cohort and the VMH cohort were analyzed separately, except for when they were directly compared and when they were combined. TDT files included timestamps of licks relayed to the TDT system through MED-PC and its lickometer, as well as the photometry streams for HisLightG and the 405 nm isosbestic control. Raw data files are available at the following link: http://doi.org/10.5281/zenodo.13981427. Data were extracted using custom Python scripts available at https://github.com/mccutcheonlab/hislightg. The photometry signal was corrected for bleaching and artefacts by using FFT-based correction method to subtract the 405 nm signal from the 465 nm signal, as previously described (Konanur et al., 2020).

Several aspects of licking behavior were analyzed: average licks per session, average number of bouts per session, and average length of bouts per session. Lick bouts were defined as a cluster of 3 or more licks with at least 10 seconds separating one bout from another. Repeated measures ANOVA with Condition (Restricted, Light, Dark, and Sucralose) as within-subjects factor were conducted for each of these parameters. We did not expect mice with PVH or VMH placements to differ in behavior. This was confirmed in a preliminary analysis finding no main effect of Region in licks, number of lick bouts, or length of lick bouts. We therefore combined mice from both regions for subsequent analyses.

Peri-event traces (hereafter, “snips”) of the photometry trace corresponding to the 5 s before a lick bout, and lasting 10 seconds after licking ended were created for each lick bout for each mouse. Of the total lick bouts, we created snips for those lasting at least 8 seconds (in the Restricted analysis of PVH vs VMH), or 4 seconds (for the analysis between conditions, due to the low number of longer lick bouts in the Sucralose condition). Artifacts were removed by comparing the absolute difference between consecutive time points and if this was greater than a threshold of 12, that lick bout was discarded. To account for different durations of each bout, only the first 6 s and last 2 s of licking were included. Snips were normalized by z-scoring, using the 5 s before licking as baseline; a baseline was calculated for each snip.

We used a strategy similar to that described in (Jean-Richard-dit-Bressel et al., 2020) to conduct waveform analyses of histamine fluctuations. First, we created a bootstrapped estimate of the mean using all lick bouts from a given region and condition, sampling 1000 times with replacement. A 95% confidence interval was then constructed around the mean. We set a consecutive threshold of 6 bins (equivalent to 0.6 s), corresponding to the low-pass filter of the photometry signal. As such, if the confidence interval did not include zero for at least 6 consecutive time bins, the period was determined as a true fluctuation in histamine. Similarly, for a comparison between PVH and VMH in the restricted condition, we compared the confidence intervals for the PVH and VMH and when they were not overlapping for at least 6 consecutive time bins, we considered them different from each other.

#### 2.4.3 Multispout experiments

For the multispout experiments, we analyzed the data with a repeated-measures ANOVA with sucrose concentration as within-subjects factor and injection (vehicle or L-histidine) as between-subjects factor. We also performed planned comparisons between injections at each sucrose concentration to test the hypothesis that at each concentration, mice with L-histidine injections would lick less than mice with vehicle injections.

## 3. Results

### 3.1 In vitro results

The fluorescent sensor HisLightG was created by combining the human H4 histamine receptor with a portion of the fluorescent sensor for dopamine, dLight1.3b (Fig 1A; Kagiampaki et al., 2023). To confirm specificity of HisLightG to histamine, we performed *in vitro* tests in which HEK293T cells were transfected and the fluorescent response to a variety of neurotransmitters was measured. Relative to control (Hank’s Buffered Saline Solution), addition of 10 µM histamine caused an approximate 150% increase in sensor fluorescence (Fig 1B; p < 0.001). Application of acetylcholine also led to a significant increase in fluorescence (p < 0.001) although this response was only a 12% increase, approximately 13 times smaller than the response to histamine, and so is unlikely to substantially contaminate fluorescent signals measured *in vivo*. All other neurotransmitters tested were without effect (Fig 1C; all p’s > 0.05).

**Figure 1.**
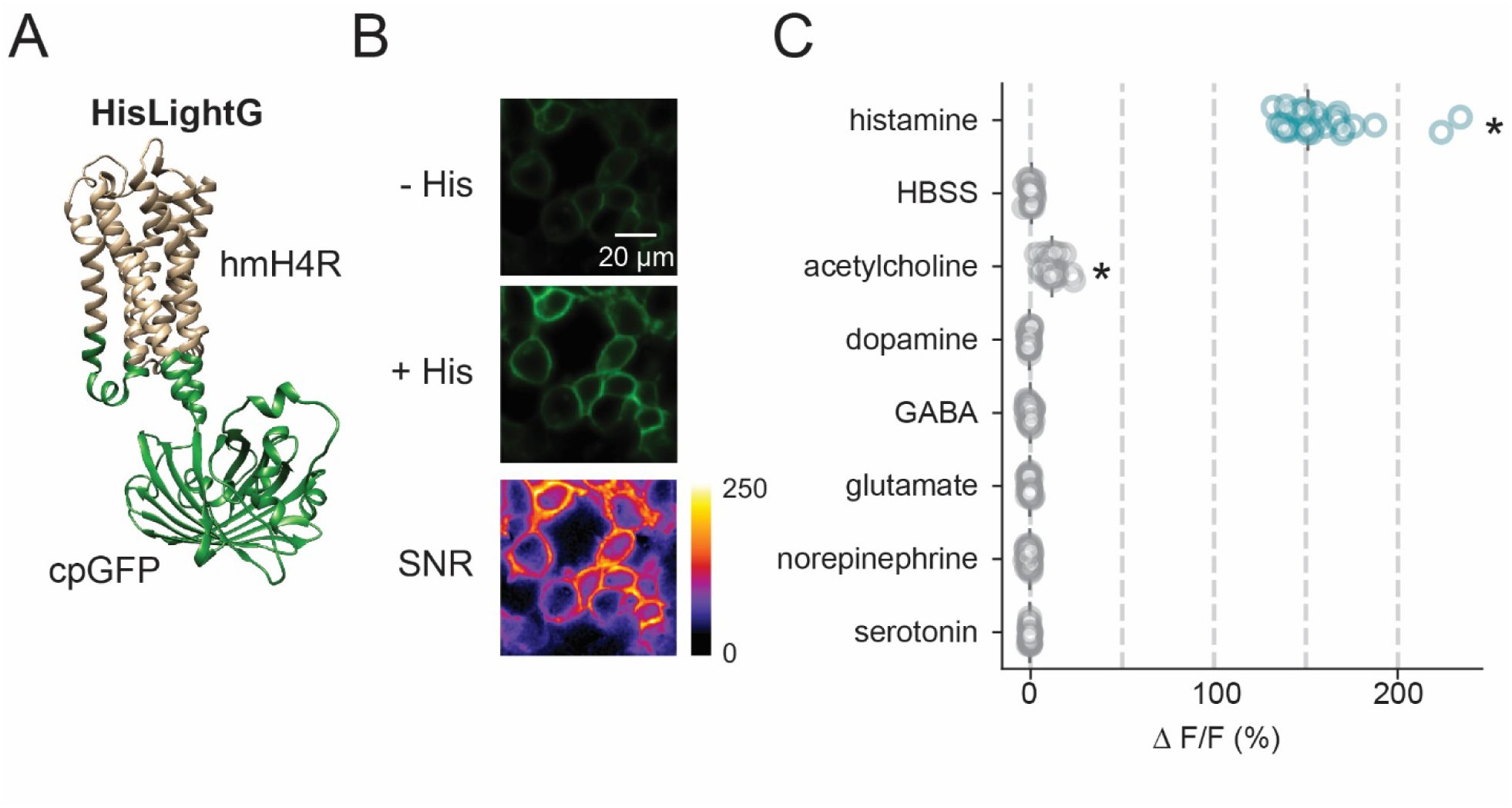
*In vitro* results. (A) Structural model of HisLightG (Kagiampaki et al., 2023) obtained through AlphaFold2 (Mirdita et al., 2022). Indicated in green are the grafted components from dLight1.3b, and in tan the original sequence from the human H4 histamine receptor. (B) Representative images of HisLightG-expressing HEK293T cells in the presence or absence of Histamine (10[μM) and corresponding signal-to-noise (SNR) heatmap. (C) Quantification of the fluorescence response of HisLightG to different non-ligand neurotransmitters (10[μM) in HEK293T cells. Data are shown as response of individual cells, with vertical lines indicating the mean[for that neurotransmitter. * p < 0.05 relative to HBSS.

### 3.2 Photometry results

#### 3.2.1 Licking behavior across time of day, hunger state, and caloric content

Licking behavior in the different conditions – Restricted, Light, Dark, and Sucralose – was compared across all mice that were included in the photometry analysis, without regard to whether the PVH or VMH was targeted. The schematic in Fig 2A details how hunger state, time of day, and nutritional value of the solution were manipulated in each of these four conditions. We plotted total lick bouts of any length (the number of bouts that included at least 3 licks which were separated by another licking bout by at least 10 seconds, with no criteria for length in time of the lick bouts) in all sessions against bout length, for each of the conditions (Fig 2B) as well as lick frequency for each condition (Fig 2C). Next, we divided these data by session and statistically examined average licks per session, average number of lick bouts per session, and average length of lick bouts per session.

**Figure 2.**
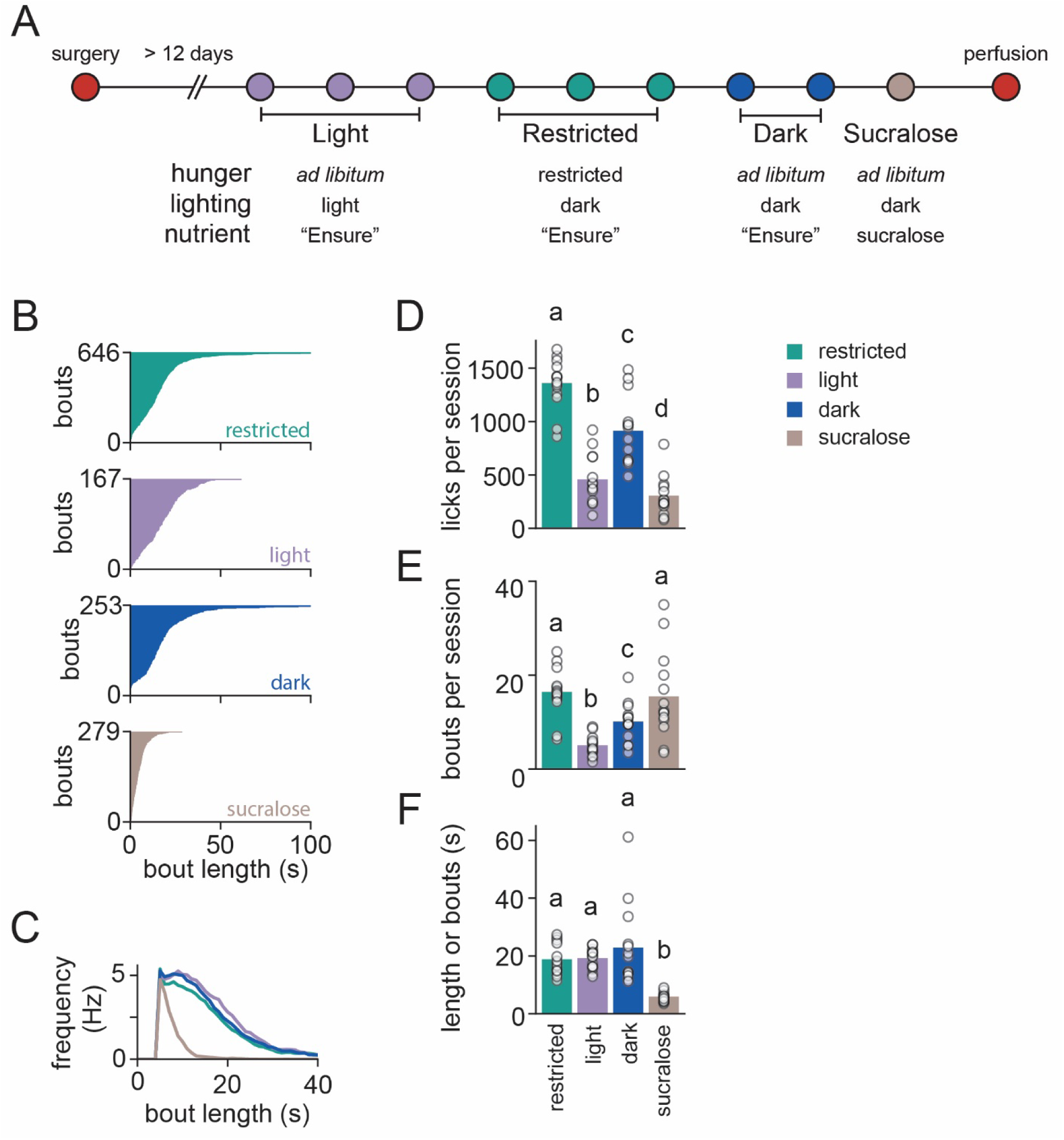
Schematic and licking behavior. A timeline of the experiment shows the number and order of photometry sessions; mice in both the PVH and VMH cohorts were tested in the same conditions and order of conditions (A). The total number of lick bouts (i.e. those that contained at least 3 licks and which were separated from another licking bout by at least 10 seconds) across all mice and sessions for each condition, plotted against bout length, is displayed (B). Lick frequency was similar in all conditions except Sucralose (C). Condition affected how much mice licked on average in each session (D). Condition also influenced the number of bouts per session (E). The average length of bouts, in seconds, also differed by Condition (F). Bars are means and circles individual data points, lower-case letters indicate p < 0.05 (e.g., a is different from b).

The average number of licks per 30 min session (Fig 2D) had a significant main effect of Condition (F_3, 39_ = 98.604, p <0.001). Post hoc tests revealed that each condition had different average numbers of licks per session. The average number of total bouts per session (Fig 2E) also differed by Condition (F_3, 39_ = 17.149, p <0.001). Post hoc tests showed that the number of bouts was equivalent in the Restricted and Sucralose conditions, which both had more bouts than the Light and Dark conditions. Dark had more bouts than Light did. Average length of lick bouts in seconds (Fig 2F) had a significant main effect of Condition (F_3, 39_ = 29.709, p <0.001). Post hoc tests showed that lick bouts were shorter in the Sucralose condition than any of the others.

Thus, by manipulating three variables - hunger state, time of day, and nutritional value of the solution - we created multiple different food intake scenarios that we could analyze to see whether each factor affected measured histamine release.

#### 3.2.2 Expression of HisLightG in the PVH and in the VMH

We examined viral spread and fiber placement in brains from mice in the PVH cohort (n=11) and the VMH cohort (n=12). In the PVH cohort 4 mice were excluded due to not having viral expression in the PVH, and in the VMH cohort 2 mice were excluded for poor virus expression and 3 for improper fiber location. This left a total of 7 mice for each region; all subsequent analyses were restricted to mice with viral expression in, and fiber tip near, the PVH (Fig 3A and Fig 3C) or VMH (Fig 3B and Fig 3D).

**Figure 3.**
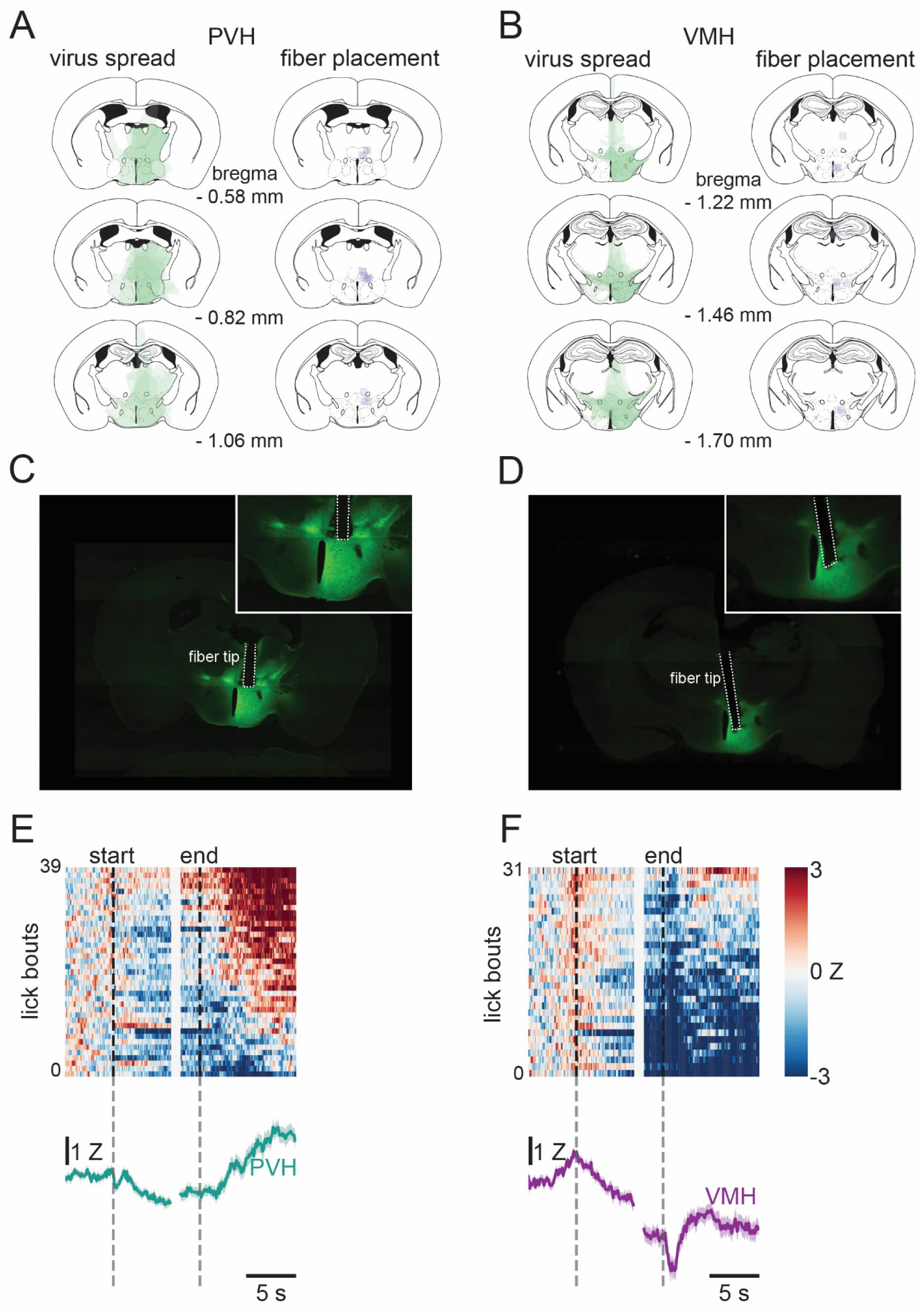
Histology and representative photometry responses in PVH and VMH. Illustration of viral spread and fiber tip location in mice included in the PVH analysis (A) and VMH analysis (B) with approximate bregma indicated. The green color shows the presence of HislightG and the grey color the fiber tip; each animal (n=7 per region) is represented by a semi-transparent shape. Images of representative animals with PVH (C) and VMH (D) injections are also shown. A heatmap showing the z-scored change in fluorescence before, during, and after a lick bout for all valid lick bouts (n=39) in a representative mouse with HisLightG in the PVH, as well as the trace of the average of those trials (E). The individual lick bouts (n=31) and the average of those for a mouse with HisLightG in the VMH (F).

#### 3.2.3 Changes in histamine associated with lick bouts, in the PVH and VMH

Our primary interest was in examining how histamine may fluctuate during and around consumption. As seen above, the most lick bouts occurred during the Restricted condition, so we looked closely at mice in this condition expressing HisLightG in the PVH and in the VMH. The z-scored change in fluorescence associated with lick bouts showed a similar pattern across lick bouts and sessions in both the representative PVH mouse (Fig 3E) and VMH mouse (Fig 3F), and it appeared that there were some differences between the regions, especially after the end of the lick bout.

To analyze these changes, we performed waveform analysis on histamine fluctuations in the PVH and VMH during the Restricted condition (Jean-Richard-dit-Bressel et al., 2020). A total of 248 lick bouts (each of the 7 mice contributing between 18-45 bouts) from the PVH and 238 lick bouts (each of the 7 mice contributing between 19-43 bouts) from the VMH were included (Fig 4A). These lick bouts were used to construct a bootstrapped mean and 95% confidence interval for each region. In the PVH, histamine was reduced below zero 1.5 s after licking started, and remained below zero for 2.8 s after licking stopped. Between 4.4 s and 10 s after licking stopped, histamine in the PVH rebounded to above zero. In the VMH, however, histamine fell below zero 0.2 s after licking started and remained suppressed throughout the lick bout and for at least 10 s after licking ceased. Comparing the confidence intervals for the histamine signal in the PVH and in the VMH revealed a significant difference (i.e., no overlap between the two regions) in the last 2 s of licking and throughout the 10 s of post-licking examined; during this time, histamine was lower in the VMH than in the PVH (Fig 4B).

**Figure 4.**
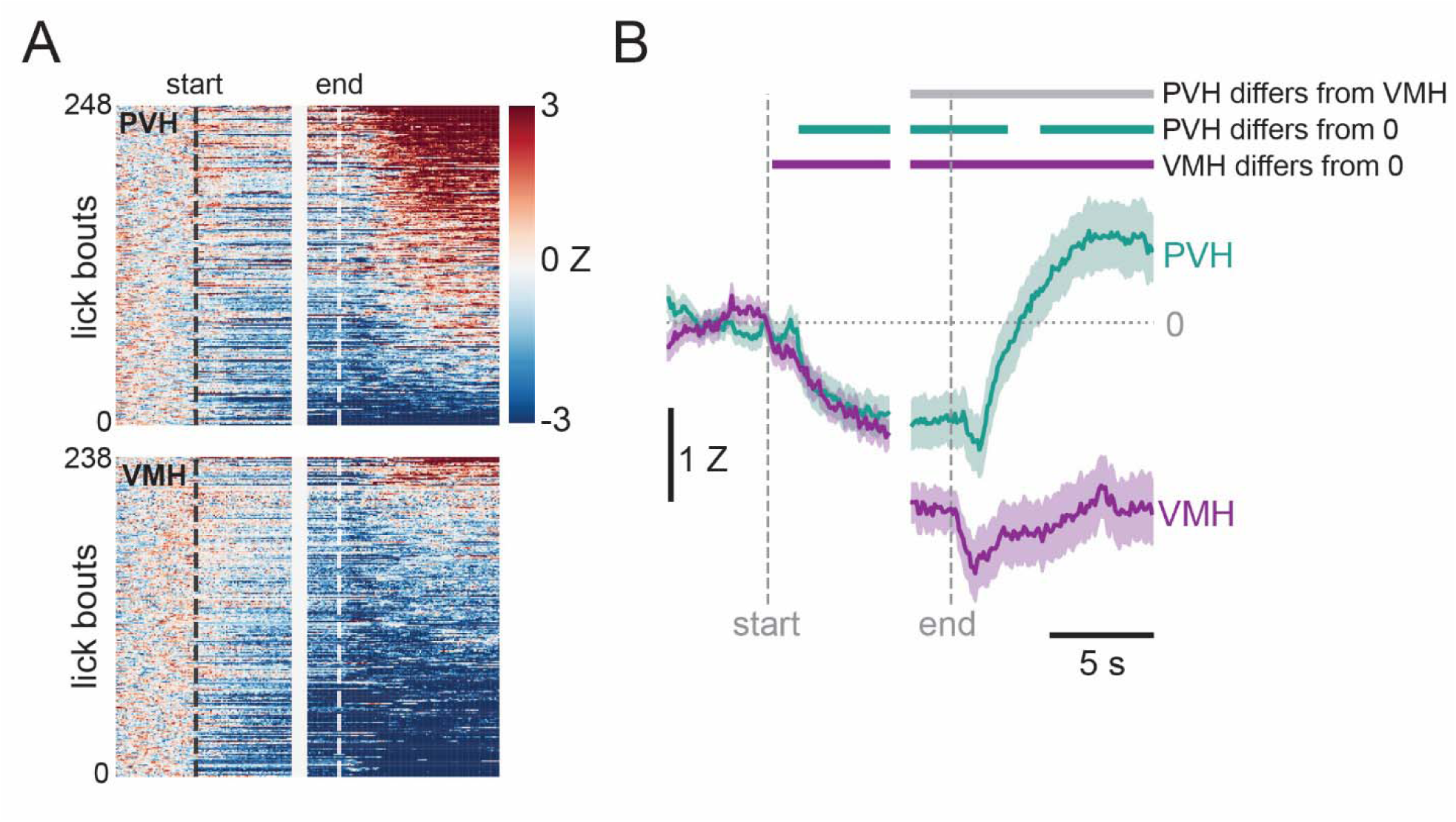
Histamine in the PVH and VMH during consumption of “Ensure” while food restricted. Each lick bout that met the criteria for photometry analysis was plotted in a heatmap, sorted based on post-licking z-score. This reveals an apparent difference between the PVH and the VMH (A). The difference in histamine was confirmed by waveform analysis, which showed that these regions differed from each other from the last 2 s of licking until at least 10 s after licking stopped; in the VMH histamine remained below baseline from the beginning of the lick bout until the end of the 10 s post-lick period, whereas in the PVH histamine was reduced during licking and for the first seconds after licking stopped, at which point it rebounded to above baseline levels (B).

In addition to the waveform analysis, we conducted a more traditional area-under-the-curve (AUC) analysis with epochs that we designated based on the behavior (i.e., the beginning and end of a lick bout, and the 10 s following a lick bout divided into two equal 5s bins). Overall, the conclusions drawn from the AUC analysis were aligned with those from with the waveform analysis (Supplemental Figure 1).

#### 3.2.3 Changes in histamine across time of day, hunger state, and caloric content

To see if histamine was sensitive to time of day, hunger state, or the caloric content of the solution, we compared the histaminergic response between conditions in the PVH (Fig 5A) and in the VMH (Fig 5B). Total lick bouts included in this analysis and the number of bouts from each mouse can be found in Table 2. In the PVH, we saw little difference in histamine in the Restricted and Dark conditions. In the Light and Sucralose condition, histamine took longer to rebound to above zero after licking stopped. In the VMH, histamine was suppressed to below zero in the Restricted, Dark, and Light conditions within a second of licking and remaining low for the full 10 s after licking stopped; in the Sucralose condition histamine did not fall below zero until the last 2 s of the lick bout.

**Figure 5.**
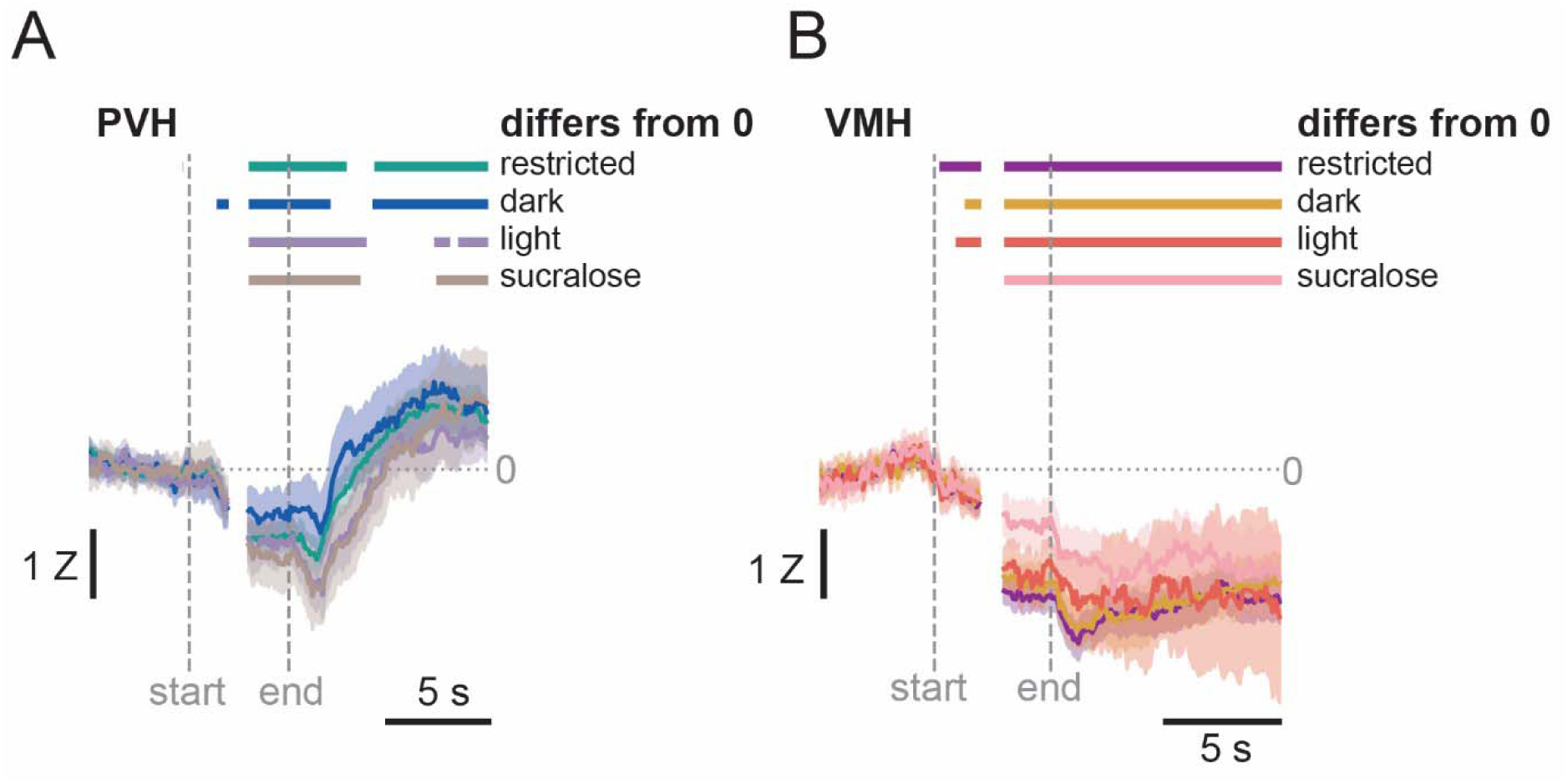
Histamine release across different conditions. In the PVH (A), histamine traces show no major differences between Dark and Restricted conditions, but a slower rebound in the post-lick period in the Light and Sucralose. In the VMH (B), there were no differences in histamine in the Restricted, Dark, and Light conditions, but no reduction in histamine during early licking in the Sucralose condition.

**Table 2:** Number of lick bouts contributing to the photometry analysis. Number of lick bouts >4 s that were used to analyze changes in histamine during different conditions. Total lick bouts included, as well as number from each of the mice, are displayed.

| Region | Condition | Total lick bouts | Bouts per mouse |
| --- | --- | --- | --- |
| PVH | Restricted | 248 | 18-54 |
|  | Dark | 107 | 5-23 |
|  | Light | 104 | 9-22 |
|  | Sucralose | 85 | 4-22 |
| VMH | Restricted | 270 | 19-54 |
|  | Dark | 126 | 10-23 |
|  | Light | 77 | 2-22 |
|  | Sucralose | 74 | 8-16 |

### 3.3 Behavioral manipulation of histamine levels

We examined licking behavior to different concentrations of sucrose in the OHRBETS multispout in mice given injections of the histamine precursor, L-histidine, or vehicle (Fig 6A). Raster plots with data from representative mice are shown in Fig 6B. These data indicate that increased licking was observed for higher concentrations of sucrose and that L-histidine suppressed this licking. Analysis of grouped data by two-way ANOVA found a main effect of injection (saline, mean 572.38 +/- SEM 64.22 vs. L-histidine, 244.88 +/- 58.09; F_1, 14_ = 14.303 p = 0.002), such that mice with L-histidine injections licked less overall than mice with vehicle injections. There was also a main effect of sucrose concentration (F_4, 56_ = 172.432, p <0.001), with mice licking more to higher concentrations of sucrose. The interaction between sucrose concentration and injection type was also significant (Fig 6C; F_4, 56_ = 11.182, p = 0.049). We performed planned comparisons between injection types at each sucrose concentration, and found that mice with L-histidine injections licked less than control mice at 0% (t_14_ = -3.882, p = 0.002), 5% (t_14_ = -2.258, p = 0.04), 10% (t_14_ = -2.243, p = 0.03), 20% (t_14_ = -4.293, p > 0.001), and 30% sucrose (t_14_ = -2.484, p = 0.026).

**Figure 6.**
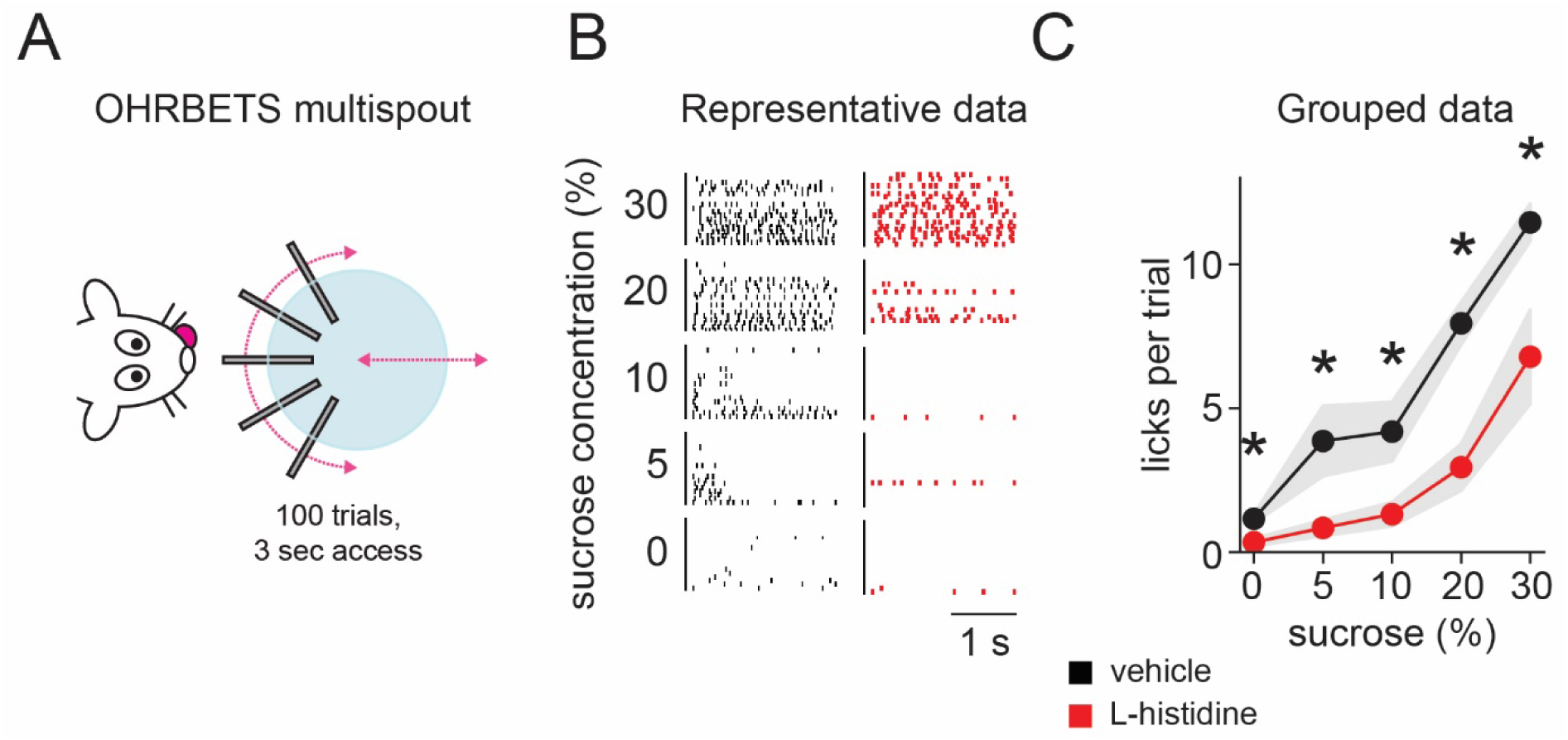
Increasing histamine suppresses sucrose intake. Mice were trained to lick in an OHRBETS multispout apparatus (A). Raster plots from representative mice show less licking in the mouse given a systemic injection of the histamine precursor, L-histidine (B), and this was also evident in the grouped data where total licks were reduced when mice were given L-histidine. Main effects of both sucrose concentration and drug administration were observed with planned comparisons showing that licks per trial were lower to at all sucrose concentrations when mice were administered L-histidine (C). Bars are means and circles individual data points. * p < 0.05

## 4. Discussion

Here, we show for the first time that a novel fluorescent histamine sensor, HisLightG, is able to detect rapid fluctuations in histamine signaling while animals engage in ingestive behavior. Although previous studies have used fiber photometry to measure histamine *in vivo* (Dong et al., 2023; Lin et al., 2023), these studies use a different sensor (GRAB-HA). Availability of multiple sensors for a single molecule is useful, as differences between sensor properties (e.g., kinetics) can make a particular sensor more suitable for a given experiment; this has been discussed in detail for dopamine sensors (Labouesse et al., 2020).

Our *in vitro* experiments clearly show that HisLightG responds to histamine. The response to histamine compared to a variety of other neurotransmitters shows a high specificity to histamine. Interestingly, acetylcholine elicited a small response, but this was an order of magnitude smaller than that of histamine, and indeed so small that it is extremely unlikely to cause any detectable signals *in vivo*. In light of this, we have a high degree of confidence that our photometry experiments were reliably measuring histamine release and not acetylcholine.

Past studies that have prevented histamine from exerting its normal function in particular brain areas have identified the VMH and PVH as the regions responsible for histamine reducing food intake (Ookuma et al., 1989, 1993; Sakata et al., 1988, 1990, 1991). Here, we examined the dynamics of histamine in these areas during eating behavior and found both similarities and differences in the histaminergic responses to consumption. In both regions, in food-restricted mice, histamine decreased as licking commenced. It remained below baseline when licking stopped. In the PVH, however, histamine rebounded to above baseline levels following the end of licking, while in the VMH it remained suppressed. The functional implications of this difference requires further investigation. A previous microdialysis experiment demonstrated a brief rise in medial hypothalamic histamine after food deprived rats were given 15 min to eat (Itoh et al., 1991). This increase in histamine was present on a timescale of minutes, indicating a modestly prolonged response of histamine to food. We show that histamine is also altered on a timescale of seconds during consumption of nutritive solutions when hungry, with changes closely linked to individual lick bouts. Interestingly, the study by Itoh et al (1991) found an increase in medial hypothalamus histamine after eating, while we found regional differences such that in the PVH histamine increased after an eating bout, but the opposite was true in the VMH.

When we examined histamine release in each region during different conditions, we again found some regional differences. In the PVH, histamine was suppressed during the lick bout in all conditions. After the lick bout, histamine rebounded in the Restricted and Dark Conditions. In the Light and Sucralose conditions, there was also a rebound but it was slower to occur. This leaves open the possibility that hunger state and caloric content of the solution influence the rebound of histamine in the PVH. Mice in the Restricted condition were hungry because of the food restriction, and in the Dark condition were presumably hungry simply from it being the time of day they normally eat. In the Light condition, the mice were likely less hungry because during the light phase they are normally sleeping, not eating, and so when these mice licked for Ensure, histamine release was different than when these same mice consumed Ensure in a different hunger state. In the Sucralose condition, the mice were tested in the dark phase and therefore equally hungry as when tested in the Dark phase, but the substance they were given to lick contained no calories. Again, this appears to have influenced histamine release in the PVH. In the VMH, in contrast, histamine remained suppressed from the start of licking until at least 10 s after licking stopped in the Restricted, Dark, and Light conditions. In the Sucralose condition, histamine showed no change from baseline during the first 2 s of licking, but was then suppressed equally as in the other conditions.

Caloric content is one possible reason why histamine responses were different in the Sucralose condition compared to the conditions in which mice were drinking Ensure, but it is also plausible that the difference was instead taste-driven. Microdialysis experiments measuring histamine in the anterior hypothalamus found that histamine levels respond to the nature of the ingested substance. Specifically, perfusates from anterior hypothalamus of food deprived rats receiving intraoral infusions of various tastants showed increased histamine after sodium chloride, hydrochloric acid, and quinine, but a decreased histamine response after saccharin or sucrose (Treesukosol et al., 2005). Interestingly, when saccharin or sucrose was paired with lithium chloride to produce a conditioned taste aversion, these tastes produced an increase in histamine (Treesukosol et al., 2005). The authors conclude that histamine release may reflect palatability of the ingested substance, with decreases indicative of high palatability. Both of our test solutions – “Ensure” and sucralose – are generally considered palatable, and this is consistent with the reduced histamine seen during licking. Whether this is related to the sustained histamine sumpression in the VMH after licking stops, and why the same solutions caused an increase after licking in the PVH, requires further study.

A fall in VMH histamine could reflect the palatability of the solution, but it is also likely that it is related to the role of histamine in suppressing food intake. Higher levels of histamine are associated with decreased food intake, so the reduction in histamine during eating could be promoting continued consumption, both within that lick bout and increasing the likelihood of subsequent lick bouts occurring due to the continued suppression. The fact that histamine in the PVH returns to baseline after a lick bout – rather than remaining suppressed as in the VMH – could mean that these regions have distinct roles in appetite regulation.

To demonstrate a role for increased histamine levels in suppressing feeding, we performed a test in a recently developed multispout apparatus (Gordon-Fennell et al., 2023) in which it was possible to test licking across several different concentrations of sucrose. In this test we found that increasing histamine levels by injecting the histamine precursor, L-histidine, suppressed licking. Thus, this links increased histamine to reduced licking and potentially satiety mechanisms. Future experiments using this method can determine whether antagonizing specific histamine receptors in specific brain regions is able to block this suppressive effect of L-histidine on licking behavior.

In summary, here we show that the fluorescent histamine sensor, HisLightG, can be used to detect rapid fluctuations in histamine release associated with eating bouts and that these patterns of release show some regional heterogeneity and sensitivity to the caloric content of the ingested substance.

## Acknowledgements

This work was supported by grants from Tromsø Research Foundation to JEM (19-SG-JMcC), the European Union’s Horizon 2020 research and innovation program (grant agreement #891959 to TP), Swiss National Science Foundation (grant #310030_196455 and #310030L_212508 to TP) and NIH to GDS (R01DA038168). The authors would like to acknowledge Rahel Möckli who assisted with histology for virus and fiber placements, and animal care staff at UiT and University of Washington.

## Author contributions

Designed research, KLV, GDS, TP, AGG-F, JEM; Performed research, KLV, AG, BB, HT, MV, AGG-F, JEM; Analyzed data, KLV, AG, JEM; Wrote the paper, KLV, JEM.

## Abbreviations

PVH: Paraventricular nucleus of the hypothalamus
VMH: Ventromedial hypothalamus
H1R: H1 receptor

---

**Figure Supplemental 1.**
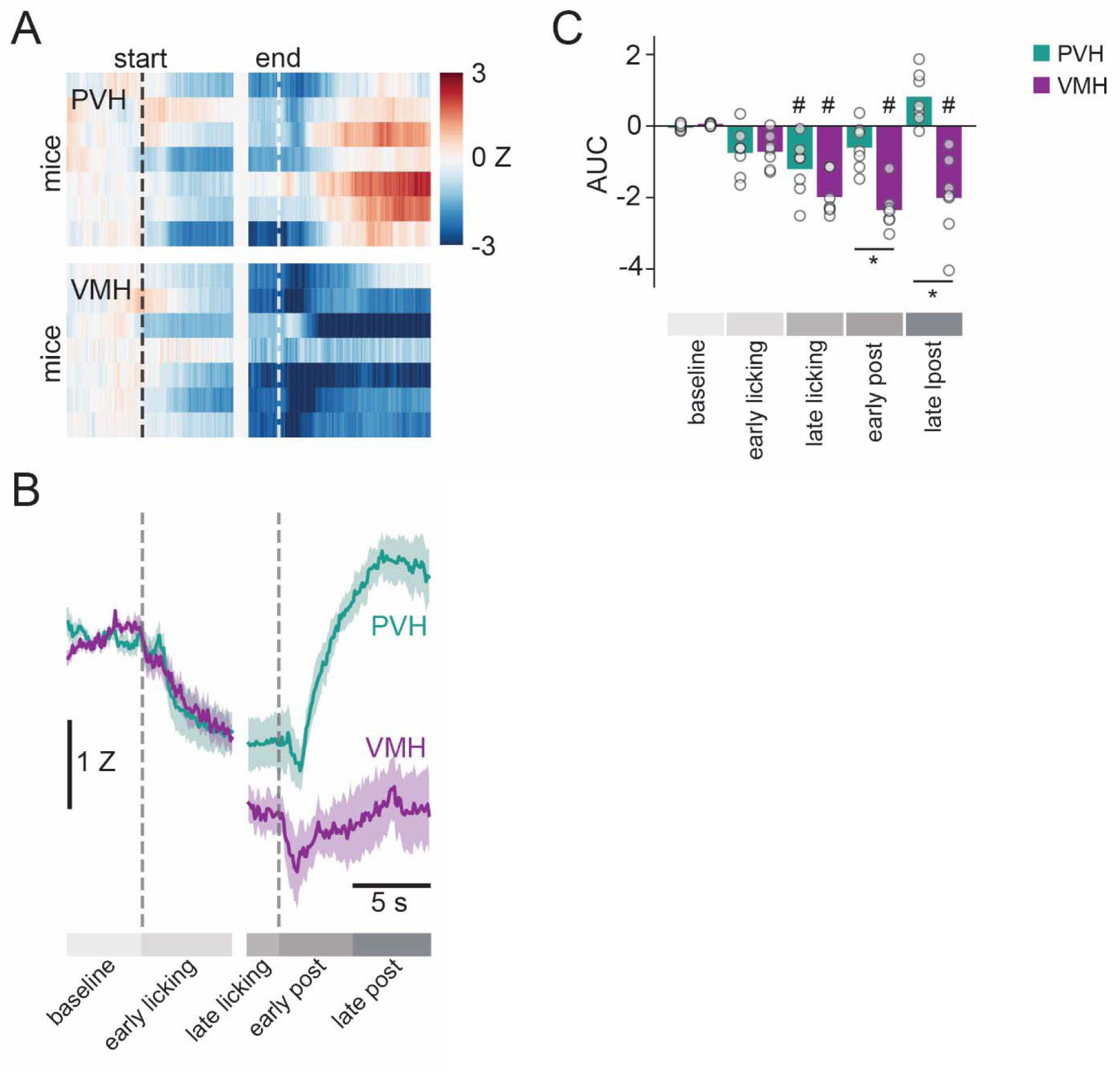
Area Under the Curve analysis of histamine in the PVH and VMH of mice drinking Ensure while food restricted. Each subject’s average Z-score across all lick bouts and sessions for mice with HisLightG in the PVH and VMH shows inter-subject consistency (A), and plotting the average for each region shows a stark difference in histamine release between regions (B). We averaged the data from all trials for each mouse (n=7 for each region) and compared changes in the AUC between regions and across time; time was divided into 5 epochs: baseline, the first 6 s of a lick bout (early licking), the last 2 s of a lick bout (late licking), the 5 s immediately after licking (early post), and the 5 s following that (late post). We found significant main effects of Time (F_4, 48_ = 21.047, p = <0.001) and Region (F_1, 12_ = 18.944, p = <0.001), as well as a significant interaction between the two ( F_4, 48_ = 18.518, p = <0.001). Holm-corrected post hoc tests revealed that in the PVH, only the late licking epoch had a lower AUC than the baseline epoch, while in the VMH, the late licking, early post, and late post epochs all had lower AUCs than baseline. Furthermore, in both post-licking epochs, the AUC of PVH and VMH differed from one another (C). Bars are means and circles individual data points, # p < 0.05 from baseline for that region, * p < 0.05 between regions for that epoch.

